# LRphase: an efficient method for assigning haplotype identity to long reads

**DOI:** 10.1101/2023.01.18.524565

**Authors:** Monica J. Holmes, Babak Mahjour, Christopher P. Castro, Gregory A. Farnum, Adam G. Diehl, Alan P. Boyle

## Abstract

**Motivation:** Understanding the functional effects of sequence variation is among the primary goals of contemporary genomics. Individual human genomes contain millions of variants which are thought to contribute to phenotypic variability and differential disease risks at the population level. However, because variants rarely act in isolation, we cannot accurately predict functional effects without first considering the potential effects of other interacting variants on the same chromosome. This information can be obtained by phasing the read data from sequencing experiments. However, no standalone tools are available to simply phase reads based on known haplotypes. Here we present LRphase: a user-friendly utility for simple phasing of long sequencing reads.

**Availability and Implementation:** LRphase is implemented in Python, and is freely available at https://github.com/Boyle-Lab/LRphase, under the MIT license. Version 1.1.0, described in this manuscript, is available through the pip and Bioconda repositories (e.g., “pip install lrphase==1.1.0”).

**Supplementary Information:** Supplementary methods are available as part of the online version of this publication.

## Introduction

Compared to the human reference genome, individual genomes differ at ∼4-5 million sites (1000 Genomes Project Consortium et al. 2015). These variants are known to affect many genomic processes (ENCODE Project Consortium 2012) and their specific effects are often affected by other nearby variants. Therefore, accurately predicting a variant’s effects requires knowledge of which other variants are present in the host genome, particularly those that are in-cis, on the same chromosomal homolog. These relationships are captured by haplotypes, which describe the set of alleles present on a single chromosomal homolog. In diploid organisms, two haplotypes exist for each chromosome, corresponding to “maternal” and “paternal” phases. We can label sequences as being “phased” if the variants it includes match the alleles for either parental haplotype. Several methods exist to reconstruct haplotypes from whole-genome sequencing (WGS) data (Bowden et al. 2019; Zhao, Liu, and Qu 2017; Li 2018; Garg, Martin, and Marschall 2016; Martin et al. 2016). While these give us the raw material needed to phase sequencing read, no currently-available tool performs simple long read phasing based on known haplotypes. LRphase provides a user-friendly, statistically-rigorous framework to address this problem.

## Description

LRphase is a command-line utility for phasing long sequencing-reads based on haplotype-resolved heterozygous variants from all contributing genomes, for example, the maternal and paternal genomes of a diploid organism (Figure 1). Long sequencing reads may be supplied as fastq files or pre-mapped reads in BAM format, while variant data are supplied in VCF format. Sequencing reads supplied as fastq are first mapped to the supplied reference genome using minimap2 (Li 2018) (Figure 1A). Mapped reads are then individually intersected with overlapping variants, and matches to each phase are counted (Figure 1B). Reads not overlapping any heterozygous variants are labeled “nonphasable”. Two modes are available for phasing reads: mode 1 uses a scoring model based on multinomial likelihoods of matching each phase; mode 2 assigns reads to the phase with the greatest number of matches to the observed genotype (Figure 1C).

**Figure 1.**
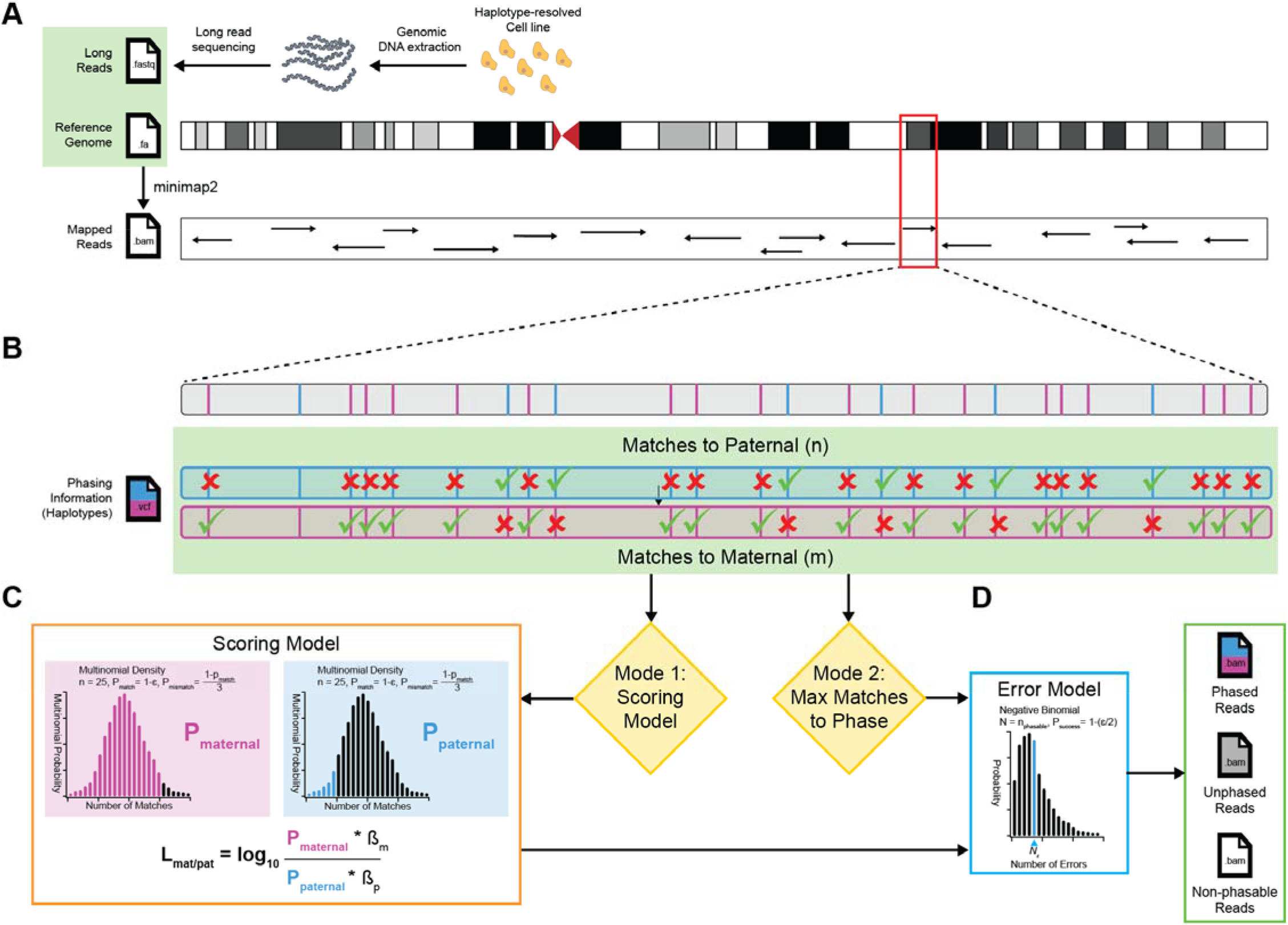
Overview of LRphase. **A)** Long sequencing reads are first prepared from genomic DNA fragments isolated from cells with available haplotype data for all parental phases. Reads are mapped to the reference genome, either within LRphase with minimap2, or externally using any desired mapping/filtering workflow, with mapped reads supplied in BAM format. **B)** Phasing begins by intersecting individual mapped reads with known phased, heterozygous single-nucleotide variants (SNVs) supplied as a VCF file. The number of matches and mismatches are counted for maternal and paternal phases (or other phase groups) and resulting counts are subsequently used to make phasing decisions using one of two scoring modes. **C)** Reads may be assigned to phases using either the scoring model (Mode 1, orange box), or by simply assigning to the phase with the greatest number of matches (Mode 2). In Mode 1, match and mismatch counts are used to calculate log-likelihoods for matching to maternal and paternal phases. Likelihoods are computed as multinomial probabilities (*P*_*maternal*_ and *P*_*paternal*_ respectively), representing the aggregate probability of obtaining the observed number of matches and mismatches to either phase given the observed sequencing error rate, □. *P*_*maternal*_ and *P*_*paternal*_ are adjusted by a Bayesian prior (ß_m_ and ß_p_) and log-likelihood ratios (LLRs) *L*_*mat/pat*_ and *L*_*pat/mat*_, are calculated to determine which phase has the most evidence for matching a given read. Reads are assigned to the phase corresponding to the greater of *L*_*mat/pat*_ and *L*_*pat/mat*_, or are labeled “unphased” if there is a tie or “nonphasable” if there is no overlap with any heterozygous variants. **D)** After all reads are phased, the false-discovery rate (FDR) is controlled by calculating the expected number of erroneous assignments, *N*_, as the mean of the negative-binomial distribution with *N* = the number of phaseable reads and *P*_*success*_ *= 1-(*□*/2)*. Phaseable reads are sorted by LLR and the *N*_□_ *\* (1-FDR)* lowest-scoring reads are relabeled as unphased. The remaining phased reads are expected to include errors at a rate corresponding to the specified FDR. Finally, labeled results are written to output file(s) in BAM format, with reads tagged to include all data used in the phasing decision.

### Scoring Model

The scoring model (Figure 1C) is fully described in the Supplementary Material. Briefly, we model the scoring distribution as multinomial, including terms for matches and mismatches to phase *i*. The □ parameter models the effect of sequencing errors on the observed numbers of matches and mismatches to phase. Thus, we parameterize the multinomial distribution: *N* = number of heterozygous sites overlapping sequence *i, P*_*match*_ = 1- □, and *P*_*mismatch*_ = □/3, since there are three mismatching nucleotides and one matching nucleotide for each variant, and calculate multinomial probabilities of matches to maternal (*P*_*maternal*_) or paternal (*P*_*paternal*_) as described (Figure 1C, Supplemental Methods). Phasing decisions are made by comparing log-likelihood ratios (LLRs) for matches to maternal and paternal haplotypes (*L*_*mat/pat*_ and *L*_*pat/mat*_). Importantly, the model may include any arbitrary combination of phase sets, not just maternal and paternal phases.

### Phasing Error Model

Since phasing decisions are based mostly on the number of matches and mismatches to each phase, we expect errors in these counts to be the primary source of phasing errors. Most counting errors can be explained by sequencing errors in the long-read data, which occur at an average rate □. At present, genotyping errors in the haplotype data are ignored as these are expected to be less-frequent and may be pre-filtered from the VCF. Sequencing errors render the observed nucleotide at position *a* uninformative. Therefore, a phasing assignment based on variant *a* reduces to a random guess between maternal and paternal phases. Assuming sequencing errors are randomly distributed among reads and occur at equal rates among heterozygous sites, each variant contributes □/2 to the expected maximum phasing error rate: (□/2)^*V*^, where *V* equals the number of variants within a sequence. In practice, we ignore the number of variants and simply use □/2 as a conservative estimate of the phasing error rate. Given these simplifying assumptions, we consider phasing decisions as Bernoulli trials with *P*_*success*_ *= 1-(*□*/2)* and model the error distribution as a negative binomial with *N* = the number of phaseable reads. To control the error rate at any given FDR, we use the mean of this distribution, *N*_, as the expected number of phasing errors, and reassign the *N*_□_ *\* (1-FDR)* lowest-scoring reads as “unphased.” Since it is independent of the scoring model, this error model has the advantage of being applicable to both of LRphase’s phasing modes.

## Discussion

Predicting a variant’s functional effects requires us to know its phase relative to other variants. For example, two non-reference SNVs within separate homologs of the same gene may elicit a compound-heterozygous loss-of-function phenotype, whereas the remaining unaffected copy may compensate when both variants are in-cis. Likewise, cis-regulatory variants likely affect only expression of the target gene copy on the same chromosomal homolog. Being able to discern between phases allows us to predict these conditions, among others.

A limitation of the current implementation is the phasing error model, which, in practice, is expected to substantially overestimate the actual number of phasing errors when multiple heterozygous sites are used in scoring. Since the probability of phasing errors decreases exponentially with increasing heterozygous site count, an improved error model would calculate and apply thresholds separately for each observed number of heterozygous sites.

LRphase has potential applications in diagnostics, clinical, and basic research into disease risk, gene regulation, and population-level genomic variation. These include targeted sequencing, phasing cut sites, high-throughput diagnostics, experimental design and hypothesis formulation, and investigation of allele-specific factor binding and/or DNA modification (methylation, etc.). Given its flexibility and ease of use, LRphase can be dropped into most analytical pipelines, making long-read phasing available to a broad audience.

## Acknowledgements

We would like to thank Carolyn Boyle and the University of Michigan Consulting for Statistics, Computing and Analytics Research (CSCAR) for providing statistical consultation in development of the scoring and error models, and members of the Boyle lab for editorial support and suggestions on the manuscript.

