## Supplemental Information for "LRphase: an efficient method for assigning haplotype identity to long reads"

### *Application Note Supplementary Information*

#### LRphase: an efficient algorithm for assigning haplotypic identity to long reads

Monica J. Holmes[2], Babak Mahjour[3], Christopher P. Castro[2], Gregory A. Farnum[2], Adam G. Diehl[2], Alan P. Boyle[1,2,†]
1. Department of Human Genetics, University of Michigan, Ann Arbor, MI, USA 48109
2. Department of Computational Medicine and Bioinformatics, University of Michigan, Ann Arbor, MI, USA 48109
3. Department of Medicinal Chemistry, University of Michigan, Ann Arbor, MI, USA 48109

**Index**

1. The LRphase Scoring Model

##### 1) The LRphase Scoring Model

The goal of LRphase is to assign all input reads to one of the known haplotypes specified in the input vcf file. Each read is assumed to have originated from a physical DNA source and the name of this genomic source (ie: the name of the cell line that the DNA submitted to sequencing was extracted from) is defined as the sample.

A vcf file can contain variants from multiple samples, and we define S as the set of all samples in the input vcf file. For a given sample *s* in a vcf file, there may be variants belonging to multiple phase sets, where a phase set is a set of phased variants that are phased relative to one another. Let $P_s$ be the set of all phase sets in sample *s* for all $s \in S$. For each phase set $p \in P_s$, let $A_p$ be the set of all phased variants specified by the vcf. The set of haplotypes for phase set *p* is defined as $H_p = \{h_1, ...h_l\}$, where *l* is the ploidy of the phase set (ie: number of distinct sequences existing in the sample population at that genomic position). For all $h_i \in H_p$, let $h_i[a]$ denote the allele for haplotype *i* at phased variant *a* for all $a \in A_p$.

Input reads must be labeled with the sample from which they originated (see methods section). Let R be the set of all input reads, $R_{mapped}$ be the set of all mapped reads in R, and for all $s \in S$ let $R_s$ be the set of all reads that originate from sample *s* in $R_{mapped}$, and let $R_{s,phasable}$ be the set of all reads in $R_s$ that overlap at least one phased variant. For each read $r \in R$, let $B_r$ be the set of all called bases for that read, let $E_r$ be the set of sequencing error rates for each called base in read *r*. For all $r \in R_{s,phasable}$, let $b_r[a]$ denote the aligned read base, and let $e_r[a]$ denote the sequencing error rate for the aligned read base at phased variant *a* for all $a \in A_p$.

| SET | TOTAL | VECTOR INFO |
|---|---|---|
| $S = \{s_1, ..., s_c\}$ | c = total samples in vcf | |
| $P_s = \{p_1, ..., p_d\}$ | d = total phase sets in a sample s | |

| | | | | | |
|---|---|---|---|---|---|
| $A_p = \{a_1, ..., a_n\}$ | n = total phased variants in phase set p | | | | |
| $H_p = \{h_1, ..., h_l\}$ | l = ploidy of haplotype for phase set p | $h_i[a]$ | Allele for haplotype i at phased variant a | | |
| $R_{s,phasable} = \{r_1, ..., r_j\}$ | j = total mapped reads for sample s with at least one phased variant overlapped | $b_r[a]$ | Read base at phased variant a | $e_r[a]$ | Sequencing error rate for read base at phased variant a |

LRphase scores follow a multinomial distribution whereby the chance of observing a match between $b_r[a]$ and $h_i[a]$ are proportional to the sequencing error rate ($\varepsilon$) in the read data. We posit that, given that we have observed a match between $b_r[a]$ and $h_i[a]$, the probability of a true match between the original sequence and $h_i[a]$ is equal to (1-$\varepsilon$). Likewise, given an observed mismatch between $b_r[a]$ and $h_i[a]$, the probability of a true mismatch between the original sequence and $h_i[a]$ is equal to $\left(\frac{\varepsilon}{3}\right)$ since there are three possible incorrect nucleotides at position *a*. Therefore, the overall probability that a read containing *n* heterozygous variants originated from haplotype $h_i$ is given by:

$$P\left(h_i\right) = \left(\frac{n!}{k_{n-m}!k_m!}\right) \times \left(\frac{\varepsilon}{3}\right)^{k_{n-m}} \times (1 - \varepsilon)^{k_m}$$

where $k_m$ is the number of variants matching $h_i$, and $k_{n-m}$ is the number of variants mismatching $h_i$ within the aligned read, and the first model term is the standard multinomial coefficient. The multinomial coefficient may be disabled via a command-line option if desired.

Multinomial probabilities are then adjusted by uniform Bayesian prior probabilities for each haplotype *i*, calculated as:

$$\beta\left(h_i\right) = \frac{1}{l}$$

For all $h_i \in H_p$. This implies an equal probability of observing any given haplotype a-priori.

The posterior probability that the observed sequence of variants matches $h_i$, then, is:

$$P(r)_{h_i} = \frac{P\left(h_i\right)\beta\left(h_i\right)}{\left[\sum_{j=1}^{l} P\left(h_j\right)\beta\left(h_j\right)\right] - P\left(h_i\right)\beta\left(h_i\right)}$$

Phasing decisions are based on the likelihood ratio of haplotype $h_i$ vs all other haplotypes in the model, calculated as:

$$LR_{h_i} = \frac{P(r)_{h_i}}{\left[\sum_{j=1}^{l} P(r)_{h_j}\right] - P(r)_{h_i}}$$

When phasing reads, LRphase calculates $LR_{h_i}$ for all haplotypes $h_i \in H_p$. Reads are assigned to the phase corresponding to max($LR_{h_1}$, ..., $LR_{h_l}$) and this value is used as the observed score for the read.
